# Clonal CD8 T cells in the leptomeninges are locally controlled and influence microglia in human neurodegeneration

**DOI:** 10.1101/2023.07.13.548931

**Authors:** Ryan Hobson, Samuel H.S. Levy, Delaney Flaherty, Harrison Xiao, Benjamin Ciener, Hasini Reddy, Chitra Singal, Andrew F. Teich, Neil A. Shneider, Elizabeth M. Bradshaw, Wassim Elyaman

## Abstract

Recent murine studies have highlighted a crucial role for the meninges in surveilling the central nervous system (CNS) and influencing CNS inflammation. However, how meningeal immunity is altered in human neurodegeneration and its potential effects on neuroinflammation is understudied. In the present study, we performed single-cell analysis of the transcriptomes and T cell receptor repertoire of 72,576 immune cells from 36 postmortem human brain and leptomeninges tissues from donors with neurodegenerative diseases including amyotrophic lateral sclerosis, Alzheimer’s disease, and Parkinson’s disease. We identified the meninges as an important site of antigen presentation and CD8 T cell activation and clonal expansion and found that T cell activation in the meninges is a requirement for infiltration into the CNS. We further found that natural killer cells have the potential to negatively regulate T cell activation locally in the meninges through direct killing and are one of many regulatory mechanisms that work to control excessive neuroinflammation.

## Introduction

The meninges are a supportive scaffold that maintains the structural integrity of the central nervous system (CNS). Composed of three main layers, the pia, leptomeninges, and dura, recent studies have identified an important role for meningeal lymphatics found in the dura in draining soluble antigens including amyloid beta that have been excreted into the cerebrospinal fluid (CSF) through the brain’s glymphatic system [1]. Likewise, impairments in meningeal lymphatic drainage exacerbate neuroimmune pathology in mouse models of traumatic brain injury [2] and amyloid pathology [3, 4]. Single-cell RNA-sequencing studies have further revealed that mouse meninges contain a unique resident macrophage population and that changes in T cell phenotype in the aging mouse dura are linked to alterations in glymphatic flux and directly impact neuroinflammation and amyloid pathology [5].

Human imaging-based studies have established impaired glymphatic flow as an important feature of Parkinson’s disease [6, 7] and increases in leukocyte density and adaptive immune signaling were observed in arachnoid granulations over the course of aging in postmortem human tissue sections [8]. Although single nucleus RNA and T cell receptor (TCR) sequencing in human meningioma and paired dura [9] as well as in Alzheimer’s disease (AD) leptomeninges [10] have provided insight into the holistic cellular composition of the human meninges, a complex view of the human leptomeninges immune landscape and its potential effects on neuroinflammation are lacking.

We performed single-cell RNA and TCR sequencing on 72,576 viable immune cells from fresh postmortem leptomeninges and brains acquired through the New York Brain Bank (NYBB) at Columbia University. We identified both myeloid and lymphoid cells across leptomeninges samples including substantial numbers of clonally expanded T cells. In addition to serving as a platform for high-dimensional identification of distinct meningeal immune subsets, samples across six neurodegenerative conditions provide insights into disease-specific alterations in meningeal immunity. We further leverage our dataset to make inferences about the activation and control of meningeal T cell activity in the meninges and its relationship to inflammatory changes in the human brain.

## Results

### Leptomeninges and brain sample collection and processing

The samples collected in this study encompassed 17 postmortem leptomeninges and 19 brains from patients diagnosed with late-onset Alzheimer’s disease (LOAD), early onset Alzheimer’s disease (EOAD), non-Alzheimer’s dementia, amyotrophic lateral sclerosis (ALS), Parkinson’s disease (PD), and frontotemporal dementia (FTD), as well as two samples from patients with no reported neurological deficits or detected neuropathologic abnormalities (**Figure 1A)**. Relevant demographic, clinical, and histopathological measures are included in **Supplementary File 1**. Briefly, leptomeninges were enzymatically digested and brain samples were mechanically dissociated, respectively. After debris removal using an in-house pipeline (**See Methods**), samples were stained with anti-CD45 BV510, anti-CD11b APC:Cy7, and propidium iodide. CD45^+^PI^-^ cells were sorted on the Sony MA900 and immediately passed through the 10x Chromium 5’ single-cell RNA-sequencing pipeline (**Figure 1A**). After strict pre-processing (**See Methods**), we retained transcriptomes from 37,491 meningeal and 35,085 brain immune cells reflecting major differences in the diversity and composition of the meningeal (**Figure 1B**) and brain parenchyma (**Figure 1C**) immunological niches.

**Figure 1:**
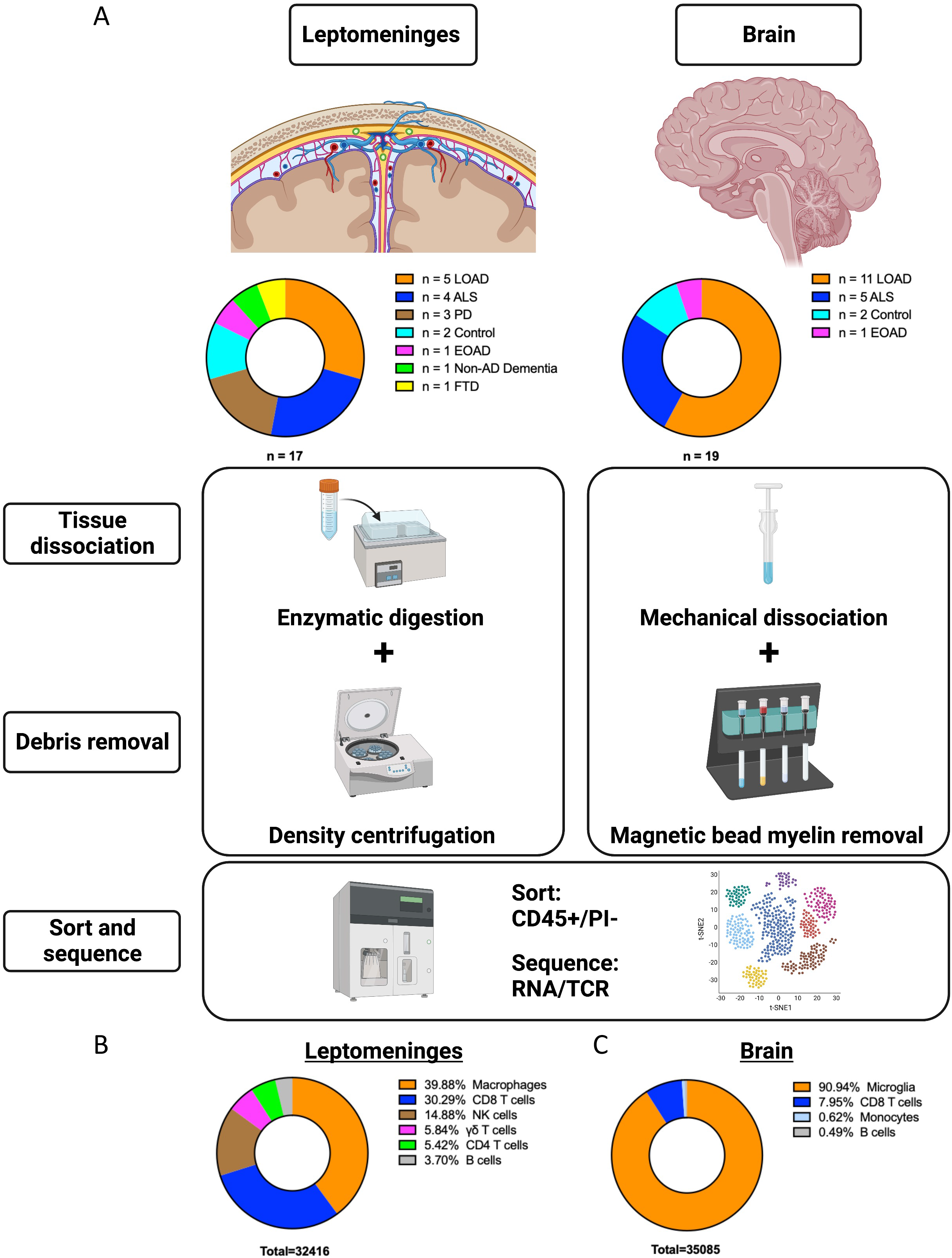
Study landscape and sample processing: (A) Breakdown of sample cohort for leptomeninges (n=17) and brain (n=19). Samples were passed through a standardized processing protocol (See Methods for details) involving physical dissociation and debris removal followed by flow sorting for live immune cells. Single cell RNA and TCR sequences were generated using the 5’ next-GEM 10X Genomics single cell sequencing protocol. Downstream clustering of 72,576 leptomeningeal immune cells across the two regions reveals a robust repertoire of adaptive and innate immune cells in the (B) leptomeninges in comparison to the (C) brain which is composed of mostly microglia.

### Resident macrophages guard the meningeal brain border

Using Seurat [11] to integrate and cluster the meningeal immune cells in our cohort, we identified 14 distinct meningeal immune clusters (**Figure 2A**). Canonical markers (**Figure 2B**) broadly identify major cell types including 4 clusters of meningeal macrophages (C1QB, LYVE1), 5 T cell clusters (CD3E), 2 NK cell clusters (NKG7), B cells (CD79B), and neutrophils (MNDA). To further characterize cell type subclusters, we used a combination of predefined markers and the FindAllMarkers function to infer potential cellular phenotypes and functions. Myeloid 1 expressed elevated levels of antigen presentation-related genes (HLA-DRA, CD74) and LOAD risk genes (APOE, TREM2, CD33) (**Figure 2C**). Indeed, Gene Ontology and Disease Ontology reflected an enrichment of mainly antigen presentation (**Figure 2D**) and neurological disease-related gene programs (**Supplementary Figure 1A**), respectively. Myeloid 4 was transcriptionally comparable to Myeloid 1 but with additional enrichment for proinflammatory cytokines (CCL3, CCL4, IL1B) (**Figure 2C**). Interestingly, in addition to the alternatively activated macrophage marker MRC1, Myeloid 3 expressed elevated levels of solute carriers SLC1A3 (GLAST) and SLC4A7 (**Figure 2C**) suggesting that these macrophages may be involved in suppressing inflammation and controlling ion and solute concentrations in the leptomeninges. Myeloid 2 was generally more transcriptionally quiescent than the other myeloid clusters but was slightly enriched for early inflammatory response genes (IER3, EGR1) (**Figure 2C**). Interestingly, Gene Ontology and Disease Ontology implicated Myeloid 2 to be involved in responding to bacterial infection (**Figure 2D; Supplementary Figure 1B**). Finally, we identified a population of Neutrophils based on their relatively low feature count [12] (**Supplementary Figure 1C)** and the expression of key neutrophil markers (S100A8, S100A9, CSF3R, MNDA) (**Figure 2C**). Meningeal myeloid cells including neutrophils egress from the skull bone marrow in response to cytokines secreted in the CSF [13] and have been implicated in the acute meningeal response to traumatic brain injury [2, 14]. Neutrophils were present in only a few patients and do not seem to correlate with clinical diagnosis (**Supplementary File 2**). However, neutrophils are highly susceptible to cellular stress and cell death induced by tissue processing and flow sorting [12], making it difficult to assess their true abundance and native transcriptional profile. Therefore, neutrophils, along with one other ambiguous cluster, were excluded from subsequent analyses. Collectively these findings suggest that myeloid cells in the meninges, especially resident meningeal macrophages, serve diverse functions that may allow them to detect and respond to infections and other harmful stimuli.

**Figure 2:**
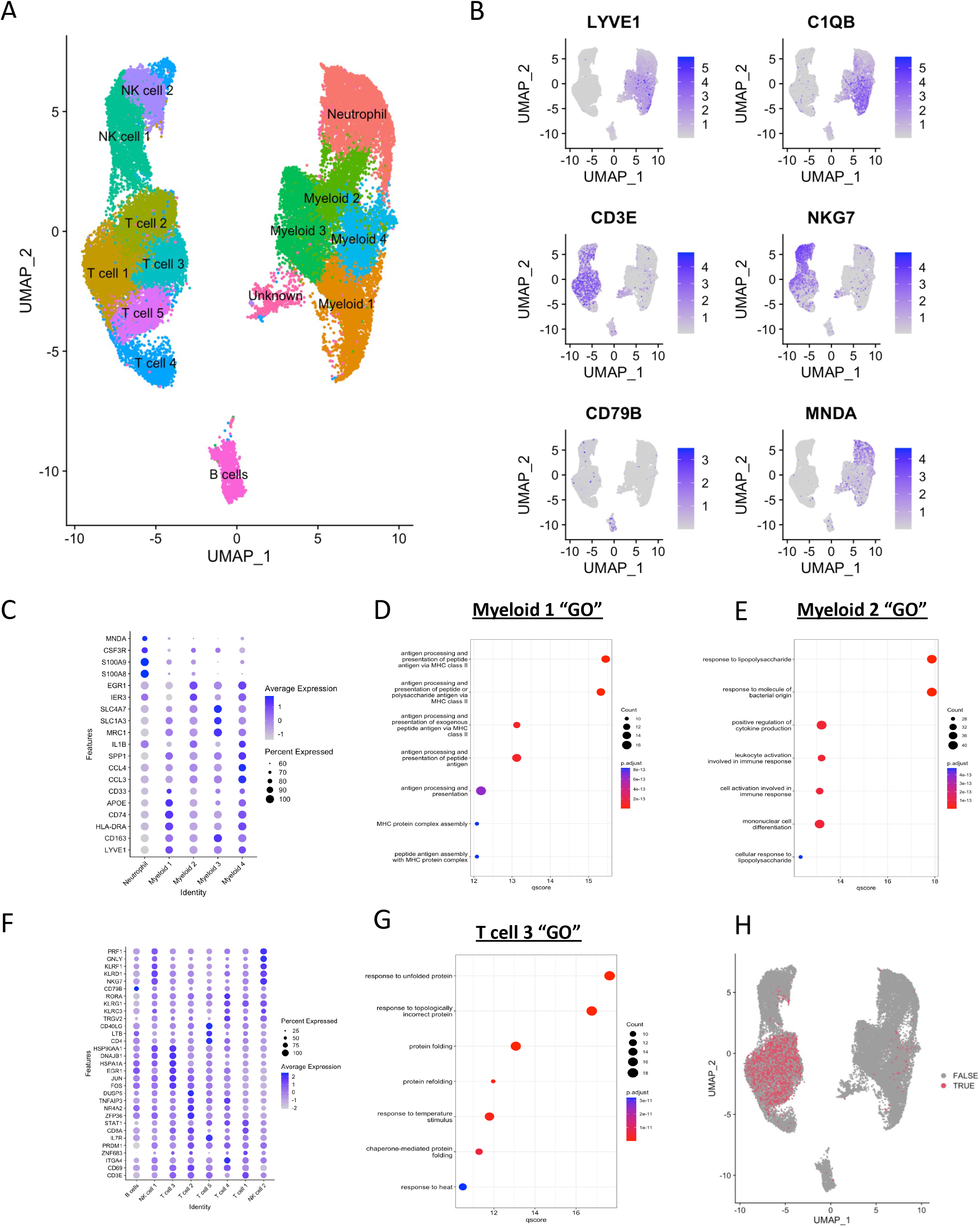
scRNA-seq identifies the leptomeninges as a diverse immunological niche. (A) UMAP depicting 14 distinct immune cell clusters identified in the human leptomeninges identified by the K-nearest neighbor clustering algorithm. (B) UMAP plots showing the expression of key marker genes in specific clusters including markers for resident meningeal macrophages (LYVE1, C1QB), T cells (CD3E), NK cells (NKG7), B cells (CD79B), and neutrophils (MNDA). (C) DotPlot of myeloid sub-cluster defining genes. Rows depict key marker genes, and columns represent distinct clusters. The blue color intensity represents the average expression of the corresponding marker in a cell cluster, while the size of the dot corresponds to the percentage of cells in that cluster that express the gene. (D) Gene Ontology analysis of key marker genes identified for Myeloid 1 by comparing the cluster’s gene expression to Myeloid cluster 2-4 and identifying upregulated genes with an adjusted p-value<0.05. (E) Gene Ontology analysis of key marker genes identified for Myeloid 2 by comparing the cluster’s gene expression to Myeloid cluster 1 and Myeloid clusters 3-4 and identifying upregulated genes with an adjusted p-value<0.05. (F) DotPlot of identified lymphoid cluster markers. Rows depict key marker genes, and columns represent distinct clusters. The blue color intensity represents the average expression of the corresponding marker in a cell cluster, while the size of the dot corresponds to the percentage of cells in that cluster that express the gene. (G) Gene Ontology analysis of key marker genes identified for T cell 3 by comparing the cluster’s gene expression to T cell clusters 1-2 and 4-5 and identifying upregulated genes with an adjusted p-value<0.05. (H) UMAP depicting alpha/beta TCR positive cells in red.

### Activated meningeal T cells patrol the brain’s borders

The remaining 6 clusters in our dataset were composed of lymphoid cells including five T cell populations with elevated levels of numerous resident memory T cell markers (CD3E, CD69, ITGA4, ZNF683, PRDM1, IL7R) (**Figure 2F**). T cell clusters 1, 2, and 3 all expressed CD8A but differed considerably in their characteristic activation and stress response characteristics. T cell 1 expressed elevated levels of the transcription factor STAT1 (**Figure 2F**) which was recently identified as a marker for early stages of memory T cell activation [15]. STAT1 is also responsible for maintaining T cell quiescence by suppressing mTOR signaling [16] which may therefore indicate that T cell 1 corresponds to a very early stage of activation. T cell cluster 2 also expressed a unique set of activation genes (ZFP36, NR4A2, TNFAIP3, DUSP5) (**Figure 2F**). Similarly, T cell cluster 3 was marked by elevated expression of acute activation markers (FOS, JUN, EGR1) in addition to various heat shock/chaperone genes (HSPA1A, DNAJB1, HSP90AA1) (**Figure 2F**). Indeed, Gene Ontology of T cell 3 marker genes implicated a relative upregulation of the unfolded protein response and response to temperature stimulus (**Figure 2G**). Interestingly, heat shock proteins are elevated at late stages of memory T cell activation [15]. Further, HSP90 can interact with alpha-4 integrins on endothelial cells to facilitate T cell adhesion and transmigration at sites of inflammation [17]. The remaining 2 T cell clusters expressed low levels of CD8A, suggesting that they may be either CD4 or non-conventional T cells. Cells in T cell cluster 5 were marked by elevated expression of CD4 and memory T cell genes (LTB, IL7R, and CD40LG) (**Figure 2F**). Interestingly, T cell cluster 4 expressed elevated levels of TCR γδ chain (TRGV2) in addition to several NK receptors (KLRC3, KLRG1) (**Figure 2F**), suggesting that they may be γδ T cells. Indeed, alpha/beta TCR sequencing confirmed that cells in T cell 4 are devoid of functional alpha/beta chains (**Figure 2H**). Meningeal γδ T cells exacerbate anxiety-like behaviors [18, 19] and short-term memory impairments in Alzheimer’s mouse models [20] by secreting cytokines including IFNγ and IL17A. Indeed, T cell cluster 4 expressed elevated levels of the transcription factor RORA (**Figure 2F**) which contributes to maintenance of T helper (Th)17-like gene programs [21]. The remaining lymphocyte populations included one cluster of B cells (CD79B) and 2 clusters of NK cells. Both NK cell 1 and 2 expressed canonical NK cell receptors (KLRD1, KLRF1) and cytotoxic proteins (GNLY, PRF1) (**Figure 2F**) with slightly higher expression of these characteristic markers in NK cell 2.

### T cells are activated locally by meningeal macrophages

Several studies have identified amyloid beta and alpha-synuclein specific T cells in the peripheral blood of patients with LOAD [22] and PD [23], respectively, and although recent reports show elevated clonal expansion of numerous TCRs in the CSF over the normal course of aging [24] and in patients with mild cognitive impairment (MCI), LOAD, and PD [25], whether clonally expanded T cells exist at the brain’s borders has not been established. Leveraging our paired RNA and TCR profiling, we found that expanded TCRs (TCRs found in two or more cells) were virtually exclusive to CD8 T cells (**Figure 3A)**. Although we identified 3 distinct clusters of CD8 T cells above, the relative levels of expansion appeared uniform across these clusters (**Supplementary Figure 2A**), reiterating that these clusters likely represent a temporal trajectory along different stages of CD8 T cell activation. As expected, CD8 T cells with hyperexpanded (5% or more of the TCR repertoire) TCRs expressed elevated levels of cytotoxicity markers (GNLY, GZMB, GZMH) in comparison to all other meningeal CD8 T cells (**Supplementary Figure 2B**). Examining the top clone in each sample revealed that the leptomeninges are a site of striking T cell clonal expansion, with 3-27% of a given patient’s overall meningeal clonal repertoire being represented by a single TCR (**Figure 3B**). We next sought to understand the effects of neurodegenerative pathology on meningeal T cell clonal expansion. Shannon entropy measures of overall clonal repertoire diversity revealed that ALS meninges tend to have more restricted repertoires in comparison to LOAD (**Supplementary Figure 2C**). Interestingly, the LOAD Shannon entropy index was comparable to that of non-neurodegenerative disease controls. Importantly, Shannon entropy did not correlate with patient sex or age (**Supplementary Figure 2D-E**), indicating that these findings are likely reflective of pathology-related effects. We next wondered whether meningeal macrophages locally influence T cell expansion in the meninges. Therefore, leveraging the inherent variability of clonal expansion in our dataset, we stratified patients based on their meningeal Shannon Entropy measures and compared the gene expression profiles of meningeal macrophages in patients with Expanded (Shannon Entropy index above the mean) versus those with Superexpanded (Shannon Entropy index below the mean) T cell repertoires (**Figure 3C**). Several inflammation and T cell homing genes including TNF, CCL3, and CCL4 were upregulated in Superexpanded meningeal myeloid cells (**Figure 3D**). Further, using both Gene Ontology and KEGG gene set enrichment to infer deregulated biological processes between these two conditions, we identified antigen processing and presentation as significantly upregulated in Superexpanded meningeal macrophages (**Figure 3E; Supplementary Figure 2F**). These data suggest that meningeal macrophages play an important role in controlling the meningeal inflammatory environment and likely engage in local antigen presentation and meningeal T cell activation.

**Figure 3:**
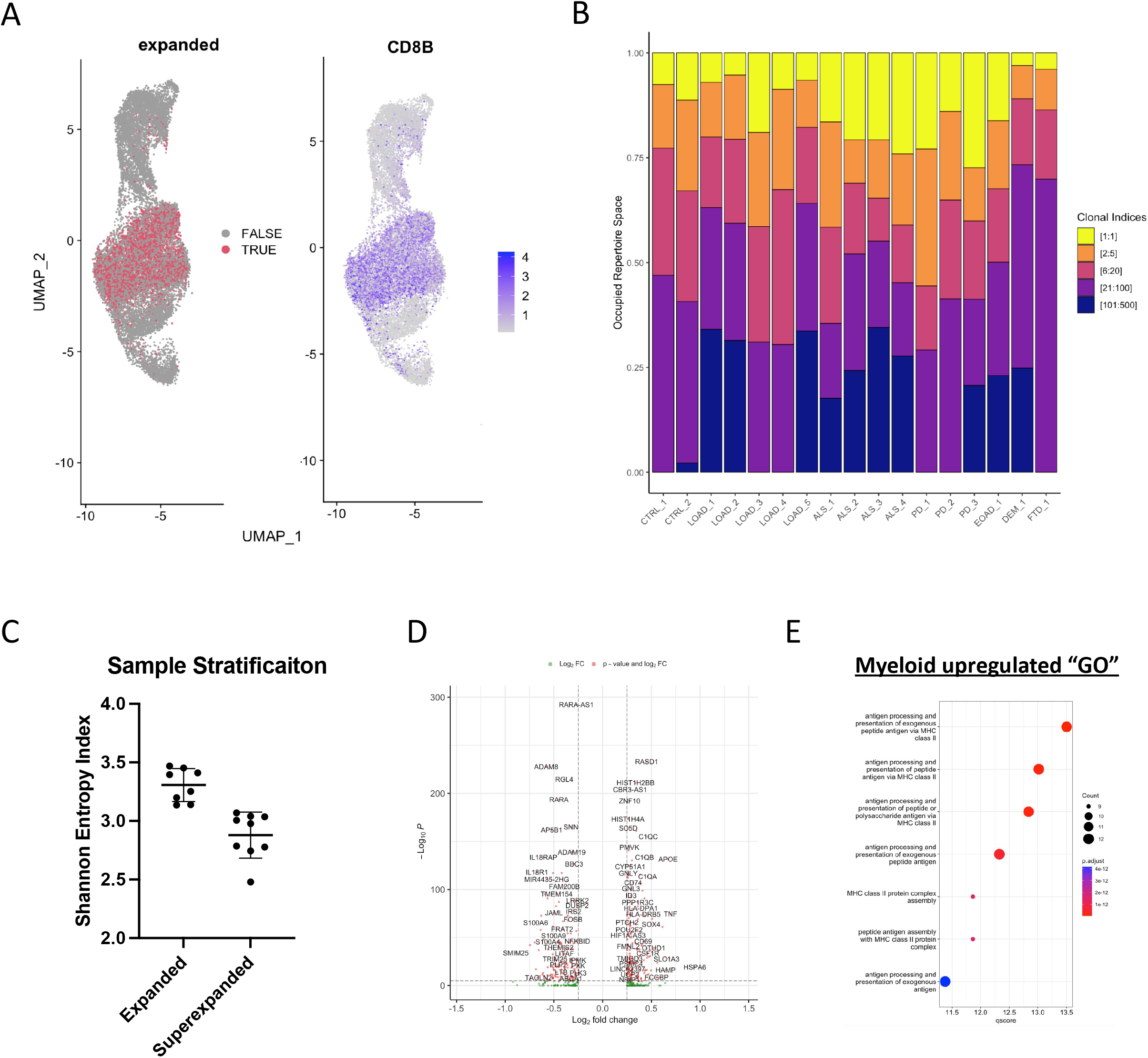
Meningeal CD8 T cells are activated locally by resident meningeal macrophages. (A) UMAPs highlighting cells with expanded (*left*) T cells defined by cells expressing a TCR found in more than one cell and cells positive for CD8B (*right*). (B) Sample breakdown of the TCR clonal repertoire. Clonal indices represent TCRs in a ranked list ordered from the TCR expressed in the highest number of cells (Clonal Index 1) to the ‘n’th most expanded TCR in the sample. (C) Shannon entropy stratification of all 17 meninges samples. “Expanded” was defined as a sample containing a TCR repertoire with a Shannon entropy index above the cohort mean, and “Superexpanded” was defined as a sample containing a TCR repertoire with a Shannon entropy index below the cohort mean. Error bars depict the SD above and below the mean in each stratified group. (D) Volcano plot of genes differentially expressed in meningeal macrophages between Superexpanded and Expanded samples. Positive Log2 fold change corresponds to genes upregulated in Superexpanded meningeal macrophages, and negative Log2 fold change corresponds to genes downregulated in Superexpanded meningeal macrophages. (E) Gene Ontology terms detected using genes identified as upregulated in Superexpanded meningeal macrophages with a p-value<0.05.

### Identification of viral antigenic triggers in the leptomeninges

Clonal expansion indicates that a T cell has been activated by its cognate antigen. Therefore, using our library of meningeal TCR’s, we sought to identify both broad neurodegenerative and disease-specific antigenic triggers of meningeal clonal expansion. Current methods to predict or identify a TCR’s cognate antigen utilize computational or targeted screening of a predefined library of antigen peptides. Fortunately, public databases including the immune epitope database (IEDB) and VDJDB compile experimentally validated TCR-antigen pairs. Although we did not identify public expanded clonotypes with exact sequence matches among meninges samples, iteratively comparing all sequences in our library using Levenshtein distance (LD) revealed several high homology matches among the CDR3 alpha and beta chains of multiple patients (**Supplementary Figure 3A**). Specifically, we identified a high homology TCR cluster between two ALS patients with reported specificity for CMV which has been associated with various neurodegenerative conditions. We also identified similar TCRs shared between a LOAD_1 (13 cells) and an ALS_1 (18 cells) patient with known specificity for influenza A (**Supplementary Figure 3A**). Interestingly, vaccination against influenza decreases risk for LOAD likely by inhibiting regulatory T cells [26] and boosting the immune response [27]. Conversely, recent reports suggest that hospitalization as a result of influenza infection significantly increases risk for multiple neurodegenerative diseases including LOAD, ALS, and PD [28]. Comparing the gene expression of these T cells to other CD8 T cells in our dataset revealed a relative upregulation of KLRC1 (**Supplementary Figure 3B**), a marker for virtual memory T cells [29]. Importantly, to ensure the T cell phenotype was conserved in clones from both samples, we performed this analysis with each patient’s clone in isolation. Indeed, KLRC1 was one of the most upregulated genes in influenza clones from LOAD_1 and ALS_1 but was only significant for ALS_1 due to the relatively low number of cells (**Supplementary File 3**). Interestingly, virtual memory T cells do not require previous engagement with their cognate antigens to confer immunologic memory to exogenous antigens and have enhanced ability to proliferate and establish resident memory T cells in response to infections including influenza [30]. Indeed, KLRC1 high cells are present throughout the meningeal CD8 T cell clusters (**Supplementary Figure 3C)** and express relatively depressed levels of the proliferation and migration suppressor RGS1 (**Supplementary Figure 3D**). Collectively, these data imply that the meninges are an important site for immune surveillance and may help protect the brain from common neurovirulent infections.

### Local control of meningeal T cell dynamics

IFNγ secreted by NK cells licensed in the gut can stimulate astrocyte-mediated killing of T cells in the mouse meninges [31]. We therefore wondered if NK cells and other immune cells in the human leptomeninges control the activation and expansion of meningeal T cells. Using Mixed-effects modeling of Associations of Single Cells (MASC) by setting expansion stratification as the contrasting feature, patient IDs as a random effect, and sex and age as fixed effect variables, we first compared the relative proportions of each cell type between Expanded and Superexpanded meninges. As expected, CD8 T cells, specifically T cell clusters 1 and 2, were more abundant in Superexpanded samples in comparison to Expanded samples (*p*= 0.001 and 0.002, respectively) (**Supplementary Figure 4A, B**). Myeloid 2, on the other hand, was relatively depleted in Superexpanded samples (p=0.02). Although not statistically significant, there was also a trend towards depletion of NK cell 1 (p=0.066) in Superexpanded meninges. With the goal of identifying specific intercellular immune signals that control T cell dynamics in the human meninges, we inferred likely ligand/receptor interactions using the CellChat R package [32]. Indeed, CellChat analysis suggested that the human leptomeninges are a site of active immune cell communication (**Figure 4A; Supplementary Figure 4D**). Among interactions between myeloid cells and CD8 T cells, we detected SPP1 (osteopontin) and GALECTIN signaling (**Supplementary Figure 4E**), two important regulators of T cell activation [33] and survival [34], respectively, suggesting that although meningeal macrophages engage in antigen presentation, they may also negatively regulate T cell dynamics in the meninges. Other intercellular interactions confirm our cell type annotations including exclusive MHC class II signaling interactions between CD4 T cells (T cell cluster 5) and meningeal antigen presenting cells (Myeloid 1, Myeloid 4, and B cells) (**Figure 4B**). As expected, MHC class I signaling was inferred as a significant signal output by all cells to varying degrees to CD8 T cells (T cell clusters 1, 2, and 3) (**Figure 4B**). Interestingly, we detected strong MHC class I signaling between NK cells and CD8 T cells (T cell clusters 1, 2, and 3) via interactions between NK cell receptor complexes (KLRK1, CD94/NKG2A, and KLRC1) and non-classical MHC class I (HLA-E) (**Supplementary Figure 4F**). Indeed, comparing gene expression in meningeal NK cells between Superexpanded and Expanded meninges revealed an upregulation of leukocyte-mediated cytotoxicity and cell killing related pathways in Superexpanded meninges (**Figure 4C**). NK cells have been shown to negatively regulate CD8 T cells by direct and indirect mechanisms in mice [35, 36], and although NK cells can directly kill PMA/ionomycin or SEB treated human CD8 T cells [37, 38], we wondered if a more physiologic T cell activation could elicit a similar effect. Indeed, we found that human NK cells kill allogeneic anti-CD3/CD28/CD2 stimulated CD8 T cells in a dose-dependent manner (*p*<0.001) (**Figure 4D**), and blocking the primary NK cell activating receptor, NKG2D (KLRK1), with a monoclonal antibody virtually abolished these effects (*p*<0.001) (**Figure 4E**). Collectively, these data suggest that NK cells can limit CD8 T cell immunity via direct contact and may regulate the degree of CD8 T cell activation, and ultimately inflammation, in the human meninges.

**Figure 4:**
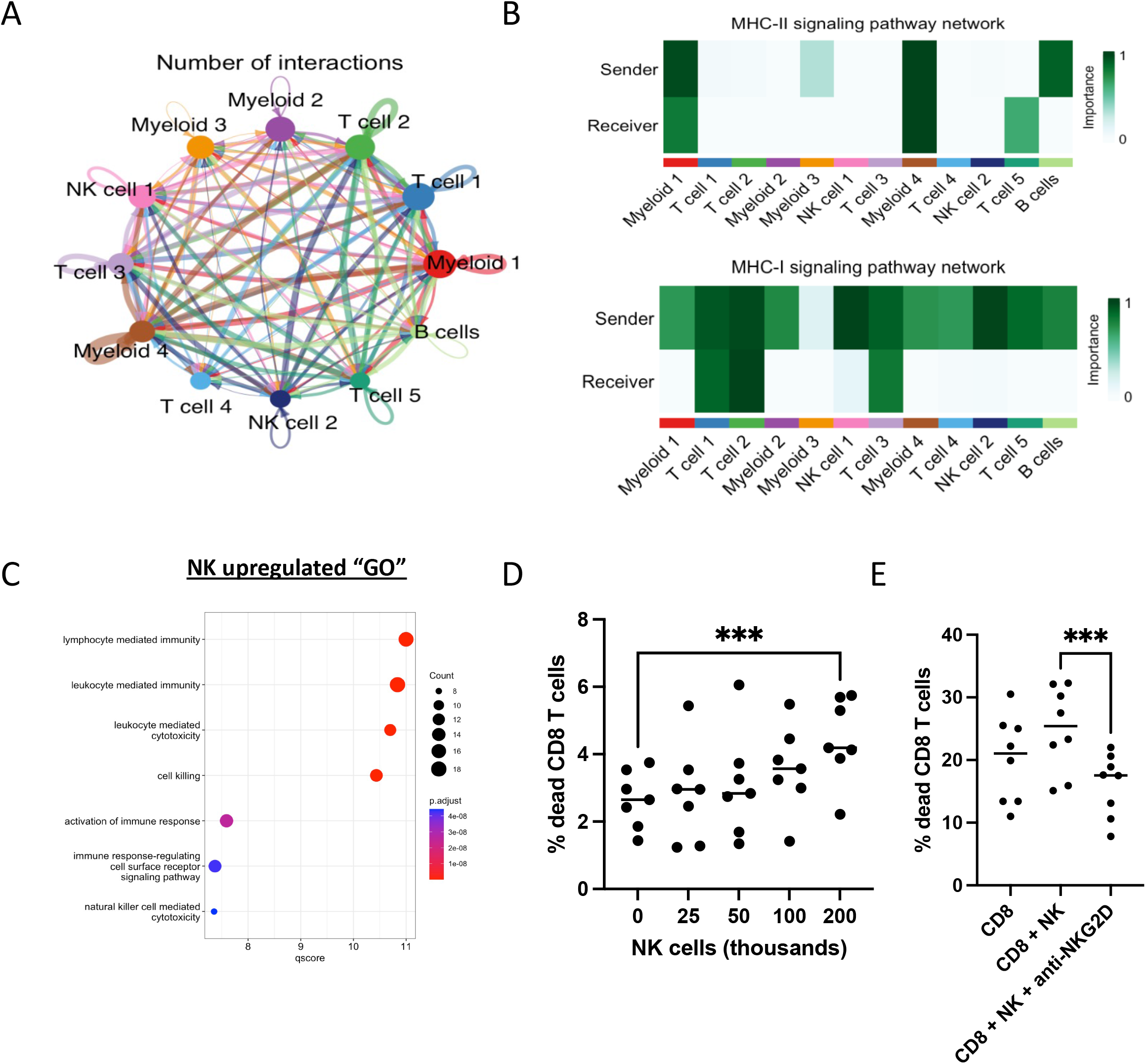
Meningeal T cell clonal expansion is regulated by intercellular immune communication networks. (A) String diagram of meningeal ligand/receptor interactions inferred using CellChat. Edge width represents the number of interactions detected. (B) Heatmap of selected intercellular interactions identified with CellChat. The intensity of green color represents the importance of a given cluster in contributing to the detected intercellular communication as a Sender and/or Receiver. (C) Gene Ontology terms detected using genes identified as upregulated in Superexpanded meningeal NK cells with a p-value<0.05. (D) The percentage of human CD8+ anti-CD3/CD28/CD2 bead stimulated cells that were dead (PI+) after 3 days in culture alone or with increasing amounts of allogeneic NK cells. All conditions n=7. ANOVA with post-hoc comparison. ***p<0.001. (E) The percentage of human CD8+ anti-CD3/CD28/CD2 bead stimulated cells that were dead (PI+) after 4 days in culture alone or with 200k allogeneic NK cells +/− anti-NKG2D neutralizing antibody. All condition n=8. ANOVA with post-hoc comparison. ***p<0.001

### Meningeal T cell activation promotes infiltration into the brain parenchyma

Meningeal immunity can influence CNS inflammation in mouse models of amyloid beta aggregation [4, 5]. To assess the effects of leptomeningeal immune changes on human brain immunity, we analyzed immune cells from 19 postmortem brain samples across 14 patients, including 10 patients overlapping with our leptomeninges sequencing cohort (**Supplementary File 1**). After strict preprocessing and integration (See Methods), we identified 13 major cell clusters (**Figure 5A**) and assigned identities based on canonical and differentially expressed marker genes (**Figure 5B)**. In addition to 9 microglia clusters, we identified 2 populations of CD8 T cells, one population of B cells, and one population of infiltrating monocytes (**Supplementary Figure 5A**). We first wondered if we could detect broad phenotypic differences between meningeal and brain immune cells. Therefore, we used RPCA to identify anchors and integrate the meninges and brain datasets into a single object to preserve expected biological differences. Differentially elevated genes included known border associated macrophage (BAM) markers including LYVE1, MRC1, and CD163 in meningeal myeloid cells and known microglia-enriched genes including TMEM119, GLDN, and OLFML3 in brain myeloid cells (**Supplementary Figure 5B**). Meningeal T cells exhibited relatively higher expression of several activation-related genes including FOS and JUN (**Supplementary Figure 5C**). However, these along with heat shock proteins are known artifacts of enzymatic digestion which was only performed on our meninges samples. Nonetheless, brain T cells were enriched for activated CD8 T cell related genes (CD8A, PFN1, CCL5) in addition to ITGA4 and CXCR3 (**Supplementary Figure 5C**) which are responsible for facilitating T cell migration across the blood brain barrier. Brain T cells also expressed elevated levels of KLRC1, potentially indicating a preference for virtual memory T cell infiltration into the brain. We next wondered whether T cells travel between the meninges and the brain. Indeed, TCRs in the two compartments overlapped, although to a variable extent (**Figure 5C**). Interestingly, patients with Superexpanded meningeal TCR repertoires exhibited significantly higher TCR repertoire overlap with the corresponding brain TCR repertoire (**Figure 5D**), suggesting that meningeal T cell activation and clonal expansion facilitates TCR-specific migration of T cells between the two compartments. Importantly, clonal expansion in the meninges did not appear to significantly affect the levels of T cell repertoire diversity in the brain (**Supplementary Figure 5D**), likely reflecting a requirement for T cells to be activated prior to brain entry. Indeed, mapping TCRs found in the brain to their corresponding cells in the meninges (**Supplementary Figure 5E**) revealed that all meninges-derived T cells in the brain are CD8 positive and have slightly depressed levels of CD27 in comparison to meningeal T cells with TCRs not found in the brain (**Supplementary Figure 5F**), suggesting that these cells may be more mature effector cells. We next wondered if TCRs found in the brain correspond to cells that are expanded in the meninges. Although singletons represented approximately 33% of all meningeal T cells (**Figure 5E**), almost all (98.47%) TCRs that were shared between the brain and meninges were expanded in the meninges (**Figure 5E**), suggesting that meningeal T cell activation and clonal expansion is a requirement for T cells to pass the blood brain barrier into the CNS.

**Figure 5:**
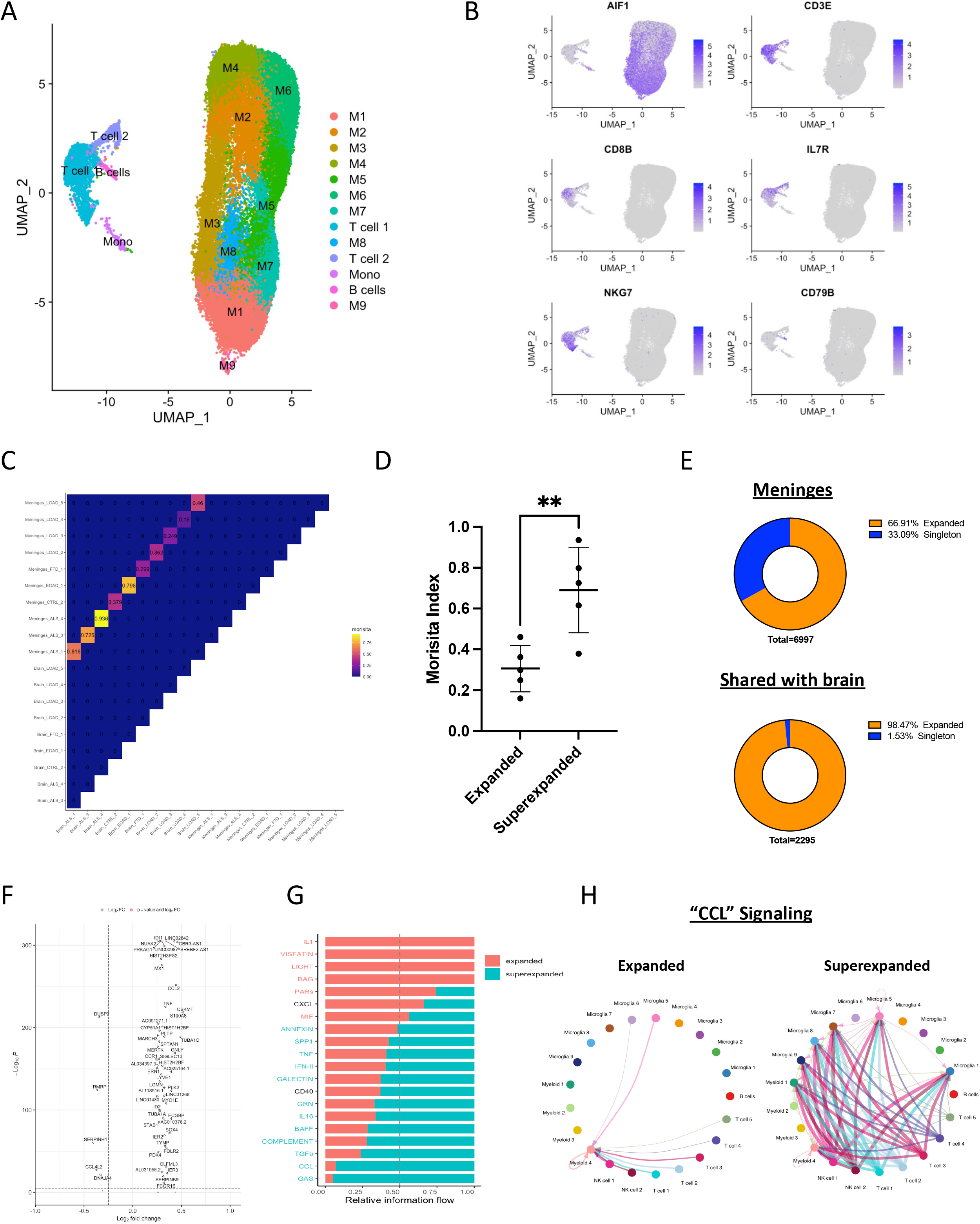
Meningeal T cell activation is a requirement for infiltration into the brain and contributes to neuroinflammation. (A) UMAP of cell clusters identified in the brain using the K-nearest neighbor algorithm. (B) UMAP highlighting cells expressing key marker genes for microglia (AIF1), Cytotoxic effector memory T cells (CD3E, CD8B, IL7R, NKG7), and B cells (CD79B). Blue color intensity corresponds to the relative level of expression of the selected gene in a cell. (C) Morisita analysis of cross-tissue TCR repertoire overlap. (D) Difference in Morisita index between patients with expanded and superexpanded meningeal TCR repertoires. Error bars denote SD around the mean. **p<0.01. (E) Donut plots illustrating the percentage of all alpha/beta TCR positive T cells in the meninges that are expanded (*top*) and all alpha/beta TCR positive T cells in the meninges that also have a TCR that is found in the brain of the same patient that are expanded (*bottom*). (F) Volcano plot of genes differentially expressed in microglia between samples with expanded and superexpanded meningeal TCR repertoires. Positive Log2 fold change corresponds to genes upregulated in Superexpanded microglia, and negative Log2 fold change corresponds to genes downregulated in Superexpanded microglia. (G) Secreted intercellular ligand/receptor interactions inferred between meningeal immune cells and microglia using CellChat. Intercellular signaling pattern names highlighted in red on the y-axis denote interactions that were inferred as significantly upregulated in expanded samples, and pattern names highlighted in blue represent interactions that were inferred as significantly upregulated in superexpanded samples. (H) String diagrams depicting differential intercellular CCL signaling between meningeal immune cells and microglia between expanded and superexpanded sample sets. Edge width represents the relative interaction strength detected between two cell clusters.

### Meningeal T cell activation influences CNS inflammation

We next wondered if meningeal T cell activation affects the inflammatory status of the brain. Comparing the proportions of brain immune clusters in patients with Expanded and Superexpanded meningeal T cell repertoires using MASC, we found that two microglia clusters, M2 and M7 were more abundant in Superexpanded meninges (*p*=0.015 and 0.013, respectively), reflecting a shift in the overall neuroinflammatory environment. Unexpectedly, we found that T cell cluster 1 is less abundant in Superexpanded meninges (*p*=0.020) (**Supplementary Figure 5G, H**), suggesting that although activated T cells migrate into the brain, infiltrating T cells are either short-lived or continuously egress from the brain, presumably through the dural lymphatics. Alternatively, it is possible that Superexpanded meninges reflect an earlier time point in pathology when T cells are beginning to infiltrate the parenchyma. Nonetheless, based on these findings, we wondered whether meningeal T cell activation affects or responds to microglial inflammation. Comparing differentially expressed genes in microglia between Expanded and Superexpanded groups revealed elevated expression of TNFα, MX1, and CCR1 in addition to CCL2 (**Figure 5F**) in microglia from patients with Superexpanded meningeal repertoires, reflecting a more inflammatory microglia state. Notably, CCL2 has been demonstrated to drive T cell recruitment to sites of CNS injury in mice [39–41]. We next used CellChat to identify potential secreted factors that facilitate communication between the meninges and brain. Among 18 significantly altered secreted signaling pathways, the CCL signaling program emerged as one of the most dynamically upregulated in Superexpanded compared to Expanded meningeal repertoires (**Figure 5G**). Specifically, whereas meningeal T cells were predicted to communicate via CCL secretion exclusively with meningeal myeloid population 4 in samples with Expanded meningeal TCR repertoires, T cells in meninges with Superexpanded TCR repertoires were predicted to communicate with numerous microglial clusters via a CCL5/CCR1 axis (**Figure 5H; Supplementary Figure 5I**). Indeed, CCL5 is elevated in ALS CSF [42], and inhibiting CCR1 was able to reduce microglial inflammation in a mouse model of intracerebral hemorrhage [43]. Notably, neutralizing CCR1 is an ongoing target of numerous clinical trials for reducing inflammation in a variety of diseases including multiple sclerosis [44]. Collectively, these data suggest that microglia and activated meningeal CD8 T cells likely interact in a bidirectional chemokine-mediated communication network and that chemokine signaling may be a potential therapeutic target for modulating meningeal T cell-mediated inflammation in the degenerating brain.

## Discussion

In this study, we comprehensively characterized the immunological landscape of the human leptomeninges and brain across multiple neurodegenerative diseases. In addition to defining the diverse immune populations in the human meninges, we established phenotypic and functional parallels between human and mouse meningeal immune dynamics. Specifically, we found that human meningeal macrophages engage in local antigen presentation and directly contribute to CD8 T cell-mediated adaptive immune surveillance of the CNS. Interestingly, we found that meningeal T cells ubiquitously express numerous resident memory T cell markers which may allow meningeal T cells to rapidly respond to potentially deleterious CNS antigens.

Although the most expanded TCRs were not annotated in publicly available TCR/peptide matching databases, we identified several expanded virus-specific TCRs in the meninges across multiple patients which is consistent with previous mouse studies showing the importance of meningeal immunity in controlling CNS viral infection [45]. Specifically, we identified TCRs in one ALS patient and one LOAD patient with significant CDR3 alpha/beta sequence homology to a previously annotated influenza-specific TCR. Further examination of the transcriptional profile of these potential influenza-specific clones revealed similarity to recently described innate-like virtual memory T cells that can rapidly mount a memory response without prior antigen exposure. Although severe influenza infection is associated with numerous neurodegenerative diseases [28], preclinical studies are necessary to determine whether meningeal innate-like CD8 T cells, specifically those with specificity for influenza contribute to neuropathology. Nonetheless, these findings reiterate a role for leptomeningeal immune surveillance in rapidly responding and potentially protecting the brain from viruses and other immunogens.

Using a combination of computational and experimental approaches, we further highlight potential regulatory mechanisms that restrain meningeal T cell clonal expansion including meningeal macrophage-derived osteopontin. In addition to serving as a regulator of microglia phagocytosis [46], osteopontin is a known immune checkpoint molecule for CD8 T cells [33]. Further studies should therefore assess the role of osteopontin in regulating meningeal macrophage antigen uptake and T cell dynamics in preclinical models of neurodegeneration. We also identified NK cells as a key altered population in Superexpanded meninges that may regulate CD8 T cells through direct killing via the NKG2D receptor. Although it is unclear if NK cells target all CD8 T cells or a specific subset, these findings could lead to cell therapies designed to modulate CD8 T cell-driven neuroinflammation.

In addition to insights into the dynamic changes that occur locally in the meninges, paired brain and meninges RNA/TCR sequencing from 10 patients allowed us to make important observations about the relationship between meningeal T cell activation and CNS immunity. Indeed, TCR sequences were shared among expanded T cells in both the brain and meninges across all samples, and in patients with more meningeal TCR expansion there was more overlap with their corresponding brain TCR repertoires. However, paradoxically we noticed an inverse correlation between meningeal T cell clonal expansion and T cell count in the brain parenchyma. Importantly, comparing the microglial signatures between Expanded and Superexpanded meningeal samples revealed an elevated level of CCL2 production by Superexpanded microglia, suggesting that the relatively low level of T cell infiltration in the Superexpanded samples may be reflective of an early phase of T cell activation preceding infiltration into the CNS rather than a defect in T cell egress out of the meninges into draining lymph nodes as seen in the aged mouse dura [5]. Indeed, preclinical studies will be necessary to assess whether microglial CCL2 recruits T cells from the meninges to the degenerating brain.

Finally, using CellChat we found that meningeal T cells likely influence microglial inflammation through the secretion of cytokines including CCL5. Interestingly, *in vitro* studies in mouse microglia suggest that CCL5 promotes pro-inflammatory signaling programs and increases microglial migration [47]. It is therefore feasible that cytokines secreted by meningeal T cells, including CCL5, promote microglia to migrate towards perivascular regions to help facilitate infiltration into the parenchyma. Indeed, recent mouse studies found that meningeal T cell-derived IFNγ decreases lymphatic drainage by altering cadherin distribution on lymphatic endothelial cells [48]. Further preclinical studies will be necessary to assess the role of meningeal T cell-derived cytokines in modulating endothelial cells and other blood-brain barrier associated cell types.

Collectively, our findings implicate the meninges as an important site of CD8 T cell activation that may be controlled by both microglial and local meningeal immune cell intercellular dynamics. Although we identified important disease-related trends pointing towards a more profound meningeal immunity disruption in ALS and PD in comparison to LOAD and controls, further studies with larger sample size are needed to assess when these changes occur over the course of disease and whether they contribute to neurodegeneration. Numerous recent studies have observed changes in the proteome and immune cell compositions of the CSF in both aging and neurodegeneration contexts [24, 25, 49], paving the way for new potential biomarkers for neurodegenerative disease progression. However, it is unclear how these peripheral changes relate to meningeal, and ultimately CNS immunity. Nonetheless, given its physiologic placement at the border between the periphery and the CNS, intercellular and chemokine-mediated dynamics in the leptomeninges could prove to be an effective avenue for biomarker development and for designing drugs to modulate CNS inflammation in certain neurodegenerative diseases.

## Methods

### Source of human brain and leptomeninges

Human postmortem brain and leptomeninges were collected by the New York Brain Bank at Columbia University using an established tissue banking pipeline [41]. After careful dissection of regions of interest, tissues were stored in Hibernate-A medium (Gibco cat. #A1237501) containing B27 serum-free supplement (Gibco cat #17504044) and stored at 4C until ready for further processing.

### Tissue dissociation and sorting

Postmortem tissue was processed immediately after collection and passed through a variety of dissociation and debris removal procedures. Leptomeninges were placed in 1mg/mL of Collagenase D (Roche cat. #51657128) dissolved in PBS and incubated at 37C for 10 minutes. Tissue was then mashed through a 70um strainer using the plunger of a 3mL syringe to achieve a single cell suspension. Cells were subsequently spun at 300g for 10 minutes and subjected to debris removal using a commercially available density gradient-based Debris Removal Solution (Miltenyi cat. #130-109-398). Postmortem brain was mechanically dissociated using a combination of mincing with a razor blade and dounce homogenization in a 15mL glass tissue homogenizer.

Dissociated tissues were passed through a 70um filter and cleared of myelin using Myelin Removal Beads (Miltenyi cat. #130-096-433) and the MultiMacs Cell24 Separator. After debris removal, leptomeninges and brain cells were pelleted at 300xg for 10 minutes and incubated with anti-CD11b APC/Cy7 (BioLegend cat. #101226) and anti-CD45 BV510 (BD Biosciences cat. #563204) antibodies at 1:100 dilution in staining buffer (2% FBS in PBS) for 20 minutes. Cells were pelleted and resuspended in staining buffer containing propidium iodide (PI) at a 1:1000 dilution. CD45^+^/PI^-^ cells were sorted on the Sony MA900 into the A1 well of a 96 well plate (Eppendorff cat. #951020401) pre-coated with 2% FBS. All sorting was performed using a 100um nozzle.

### 10x Genomics Chromium single cell 5’ library construction

Cell capture, amplification, and library construction was performed on the 10x Chromium platform using the manufacturer’s publicly available protocol. Briefly, after sorting a fraction of cell suspension was used to estimate total cell count and viability. All remaining sorted cells were loaded in the Chromium platform using the 10x Genomics Next GEM Single Cell 5’ v2 for scRNA and TCR-seq. Libraries were prepared using 10x library reagents in accordance with their 5’ RNA and TCR sequencing library protocol and sequenced on the NovaSeq 6000. RNA and TCR reads were aligned to the Hg38 genome and clonotype/contig matrices were generated using Cell Ranger (v6.1.2).

### Pre-processing of paired scRNA-seq and scTCR-seq data

Sequencing data analyses were conducted using R (v4.0.0 for data filtering and sample integration, v4.2.2 for downstream analyses). Barcoded matrices from CellRanger were processed in Seurat (v4.3.0) by retaining features expressed in at least 3 cells. Low quality and dying cells were removed by filtering out cells with fewer than 200 features and greater than 10% of reads aligning to mitochondrial genes, respectively.

### Sample integration and cell cluster annotation

Leptomeninges Seurat objects were integrated first by identifying anchor features using Canonical Correlation Analysis (CCA) with the FindIntegrationAnchors function in Seurat. Clustering was then performed using the K-nearest neighbor algorithm utilizing 30 dimensions and a resolution of 0.6. Cluster identities were defined using both accepted canonical markers and genes identified using the FindAllMarkers function. KEGG GSEA analysis was performed using the gage R package by inputting all differentially expressed genes. Gene Ontology and Disease Ontology analyses were performed using the clusterProfiler R package by selecting differentially expressed genes with a *p*-value<0.05. Brain single cell data was similarly integrated as above and clustered using 10 dimensions and a resolution of 0.5. Cross-tissue single cell comparisons were performed by integrating the Seurat objects generated above using reciprocal PCA (RPCA) and clustering using K-nearest neighbor with 30 dimensions and a resolution of 0.5. Cluster proportions were compared across conditions using Mixed-effects modeling of Associations of Single Cells (MASC) [50] by setting expansion stratification as the contrasting feature, the patient IDs as a random effect, and sex and age as fixed effect variables.

### TCR sequencing analysis

TCR sequences were analyzed and integrated into the single cell RNA Seurat object using the R package scRepertoire. Only TCR sequences containing both a complete CDR3 alpha and beta chain were considered for downstream clonality analyses. TCR data with more than one CDR3 alpha or beta chain detected in a single cell were excluded. TCR clones were considered identical if both the VDJC gene and CDR3 nucleotide sequences for both the alpha and beta chain were identical. Shannon entropy was measured using the clonalDiversity function in scRepertoire. Sample stratification by expansion level was performed by calculating the mean Shannon entropy across the 17 leptomeninges samples and segregating by samples with indices above the mean (Expanded) and below the mean (Superexpanded). Levenshtein distance measurements were performed using the base R function adist. TCRs were first filtered to include only those expressed in 5 or more cells and barcoded with a unique string tag (“_a”, “_b”, etc) to identify the sample source in downstream analysis and visualization.

### Inference and analysis of intercellular communication using CellChat

Ligand/receptor interactions were inferred using the CellChat package in R using standard settings. Cross-tissue CellChat analysis was performed by first subsetting the microglia from the brain sequencing Seurat object and merging with the leptomeninges Seurat object using the merge function. The resulting Seurat object was then split using the subset function to generate separate Seurat objects for Expanded and Superexpanded conditions and passed through the CellChat compare pipeline.

### NK cell killing assay

CD8 T cells and allogeneic NK cells were isolated from PBMCs from the New York Blood Center (NYBC) using the Miltenyi MACS Human CD8 T cell isolation kit (cat. #130-096-495) and the Miltenyi MACS human NK cell isolation kit (cat. #130-092-657). CD8 T cells were activated with immunoCult Human CD3/CD28/CD2 T Cell Activator beads (cat. #10970) in co-culture with NK cells in the presence of IL2 (2ng/mL) (R&D Systems cat. #BT-002) and IL15 (0.1ng/mL) (R&D Systems cat. #BT-015). NKG2D neutralization was performed by first pre-incubating 200k NK cells with 100ug/mL anti-Human CD314 (NKG2D) antibody (Invitrogen cat. #14-5878-82). Cells were spun down at 300xg for 10 minutes after which the antibody concentration was diluted to 50ug/mL for the duration of the 4-day co-culture with anti-CD3/CD28/CD2 activation beads and 100k CD8 T cells in the presence of IL2 (2ng/mL) and IL15 (0.1ng/mL).

## Data availability

Raw and analyzed sequencing data in this study will be deposited in the NCBI GEO database at the time of publication. Single cell RNA and TCR sequencing analyses will also be made available via an interactive Shiny app.

## Acknowledgments

We thank the individuals and their families who have generously donated the brain and leptomeninges tissues used in this project through the New York Brain Bank (NYBB). We thank Vilas Menon for advice on computational analyses, Michael Kissner for technical assistance and resources for sorting and single cell sequencing protocols, and the Columbia Genome Center for providing library generation and sequencing services. The work was supported by grants from the National Institute of Aging (5R01AG067581 and 3R01AG067581-04S1) and the Parkinson’s Foundation (PF-IMP-870699). This research was funded in part through the NIH/NCI Cancer Center Support Grant P30CA013696 and used the Genomics and High Throughput Screening Shared Resource. This publication was supported by the National Center for Advancing Translational Sciences, National Institutes of Health, through Grant Number UL1TR001873. The content is solely the responsibility of the authors and does not necessarily represent the official views of the NIH.

## Author contributions

R.H. and W.E conceived the study. R.H. performed the experiments. R.H., S.L, X.F., B.C., D.F., and H.X. provided technical assistance. R.H., C.S., E.B., N.A.S, and W.E provided intellectual input. A.F.T. contributed the human brain specimens. R.H. and W.E wrote the manuscript, and all authors edited it.

## Supplementary Figure Legends

**Supplementary Figure 1:**
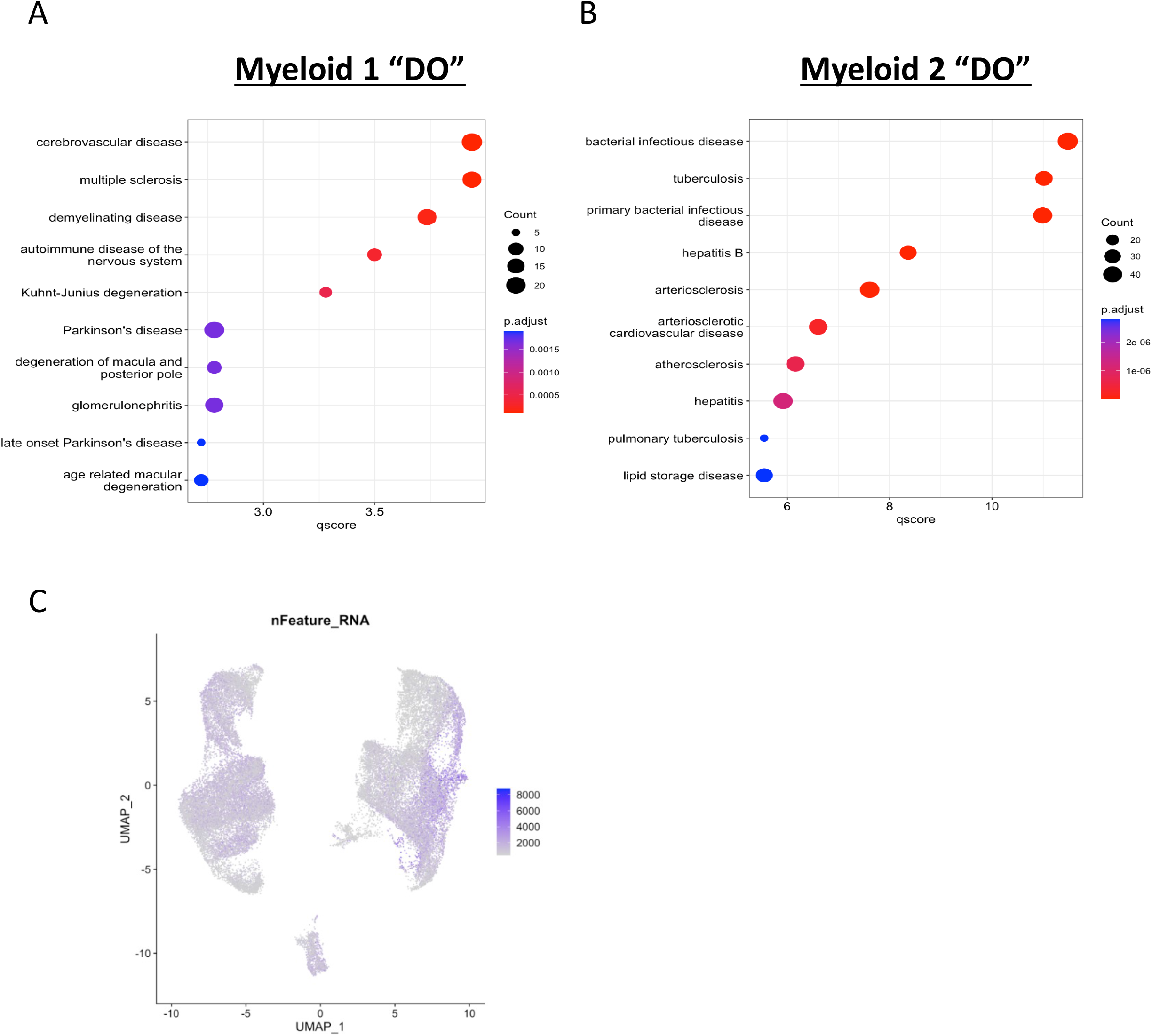
Disease Ontology of meningeal clusters (A) Myeloid 1 and (B) Myeloid 2. Gene ontology was calculated using key marker genes identified for each myeloid cluster by comparing the cluster’s gene expression to all other meningeal macrophage clusters and identifying upregulated genes with an adjusted p-value<0.05. (C) Feature plot depicting the number of genes detected in each cell.

**Supplementary Figure 2:**
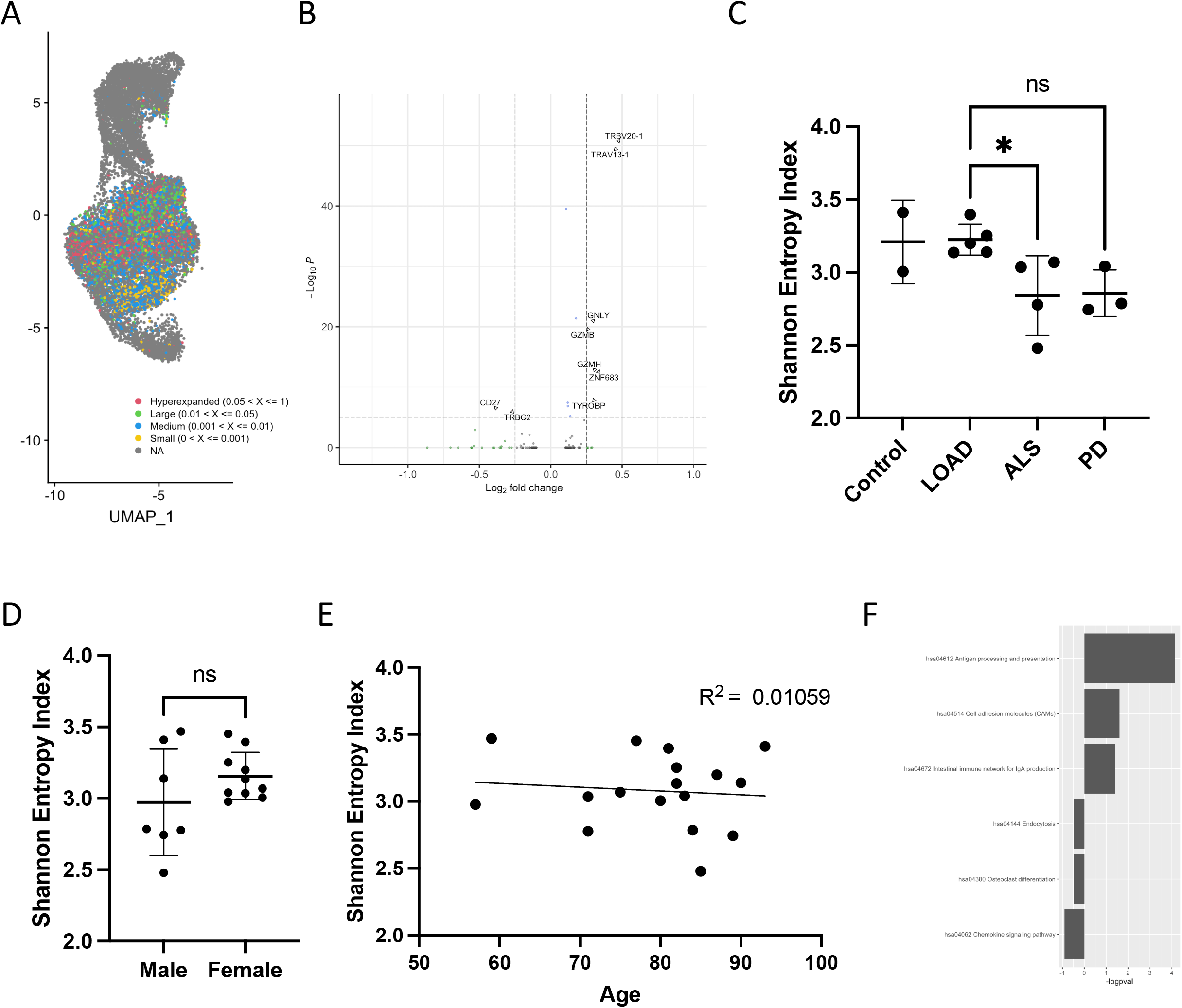
Local meningeal CD8 T cell expansion is a pathology-driven phenomenon. A) UMAP highlighting the level of TCR expansion in meningeal T cells. B) Volcano plot of genes differentially expressed between hyperexpanded and all other CD8 T cells with a full alpha/beta TCR sequence. Positive Log2 fold change corresponds to genes upregulated in hyperexpanded meningeal CD8 T cells, and negative Log2 fold change corresponds to genes downregulated in hyperexpanded meningeal CD8 T cells. (C) Shannon entropy index measure of sample TCR diversity stratified by disease. Error bars depict the SD above and below the mean in each stratified group. ANOVA with post-hoc comparison. *p<0.05. (D) Shannon entropy index stratified by patient sex. t-test. (E) Shannon entropy plotted as a function of patient age. Best fit line plotted using linear model. (F) KEGG GSEA pathways differentially represented in superexpanded versus expanded meningeal macrophages calculated by using all genes with a detected logFC>0. Positive logpval denotes pathways upregulated in superexpanded meningeal macrophages, and negative logpval denotes pathways downregulated in superexpanded meningeal macrophages.

**Supplementary Figure 3:**
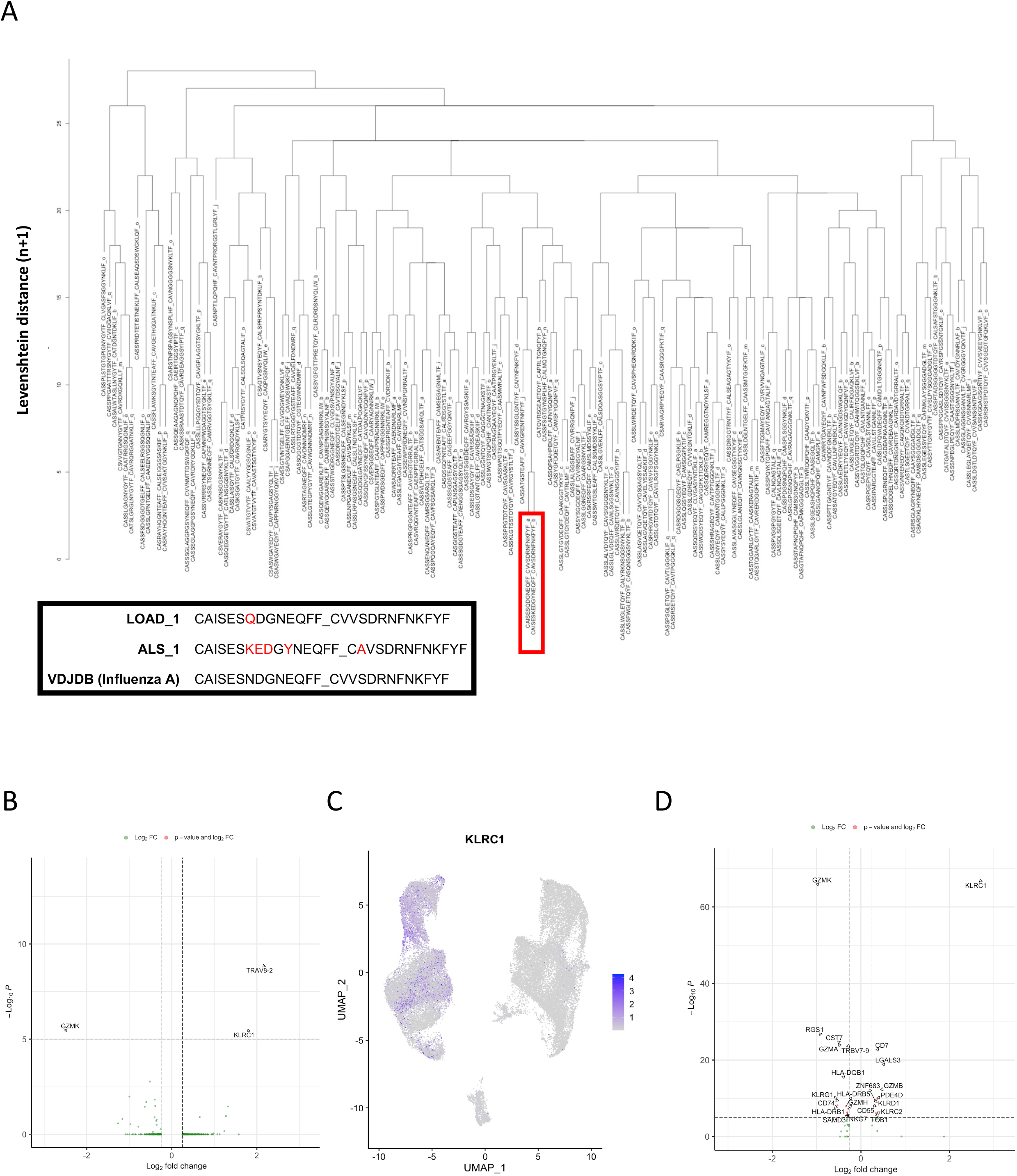
A subset of expanded meningeal CD8 T cells may be virus-targeting virtual memory T cells. (A) Dendrogram of Levenshtein distance for all meningeal TCRs expressed in 5 or more cells. TCRs were given a patient-specific barcode (ie “_a”, “_b”) to facilitate identifying the sample source of a given TCR. Red box around Levenshtein clustered TCRs and block box in lower left highlight TCRs with high homology between a patient with LOAD and another with ALS that according to a similar TCR in VDJDB may target Influenza A. (B) Volcano plot of genes differentially expressed between the shared influenza-specific clones and all other meningeal CD8 T cells. Positive Log2 fold change corresponds to genes upregulated in cells expressing the influenza-specific TCRs, and negative Log2 fold change corresponds to genes downregulated in cells expressing the influenza-specific TCRs. (C) UMAP plot showing the expression of KLRC1 across meningeal cell clusters. (D) Volcano plot of genes differentially expressed between virtual memory (KLRC1 positive, alpha/beta TCR positive) CD8 T cells and all other meningeal CD8 T cells. Positive Log2 fold change corresponds to genes upregulated in virtual memory T cells, and negative Log2 fold change corresponds to genes downregulated in virtual memory T cells.

**Supplemental Figure 4:**
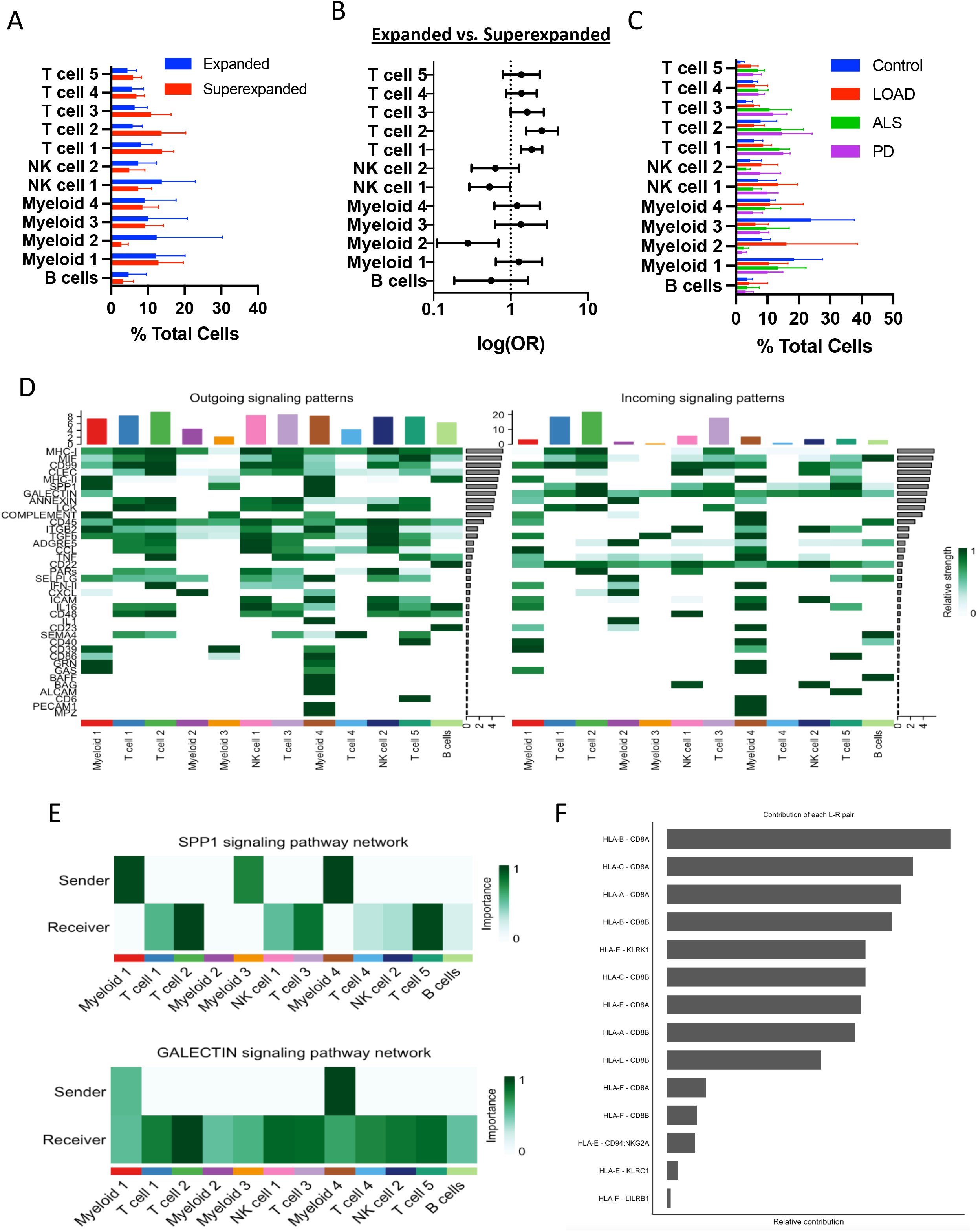
Meningeal CD8 T cell clonal expansion may be affected by local changes in key regulatory cell clusters. A) Percentage composition of cell clusters in expanded versus superexpanded meninges. Error bars correspond to 1 SD above and below the mean. B) MASC analysis of cell cluster proportional changes between expanded and superexpanded meninges. Odds ratio greater than 1 corresponds to an increased abundance of a particular cell cluster in the superexpanded group. Error bars correspond to 95% CI. C) Percentage composition of cell clusters by disease. Error bars correspond to 1 SD above and below the mean. D) Heatmap of all outgoing (*left*) and incoming (*right*) ligand/receptor signals detected between meningeal cell clusters using CellChat. E) Heatmap of selected signaling programs and the corresponding cluster-specific signals contributing to the interaction. F) Relative contribution of specific ligand/receptor pairs to the MHC-I CellChat signal.

**Supplementary Figure 5:**
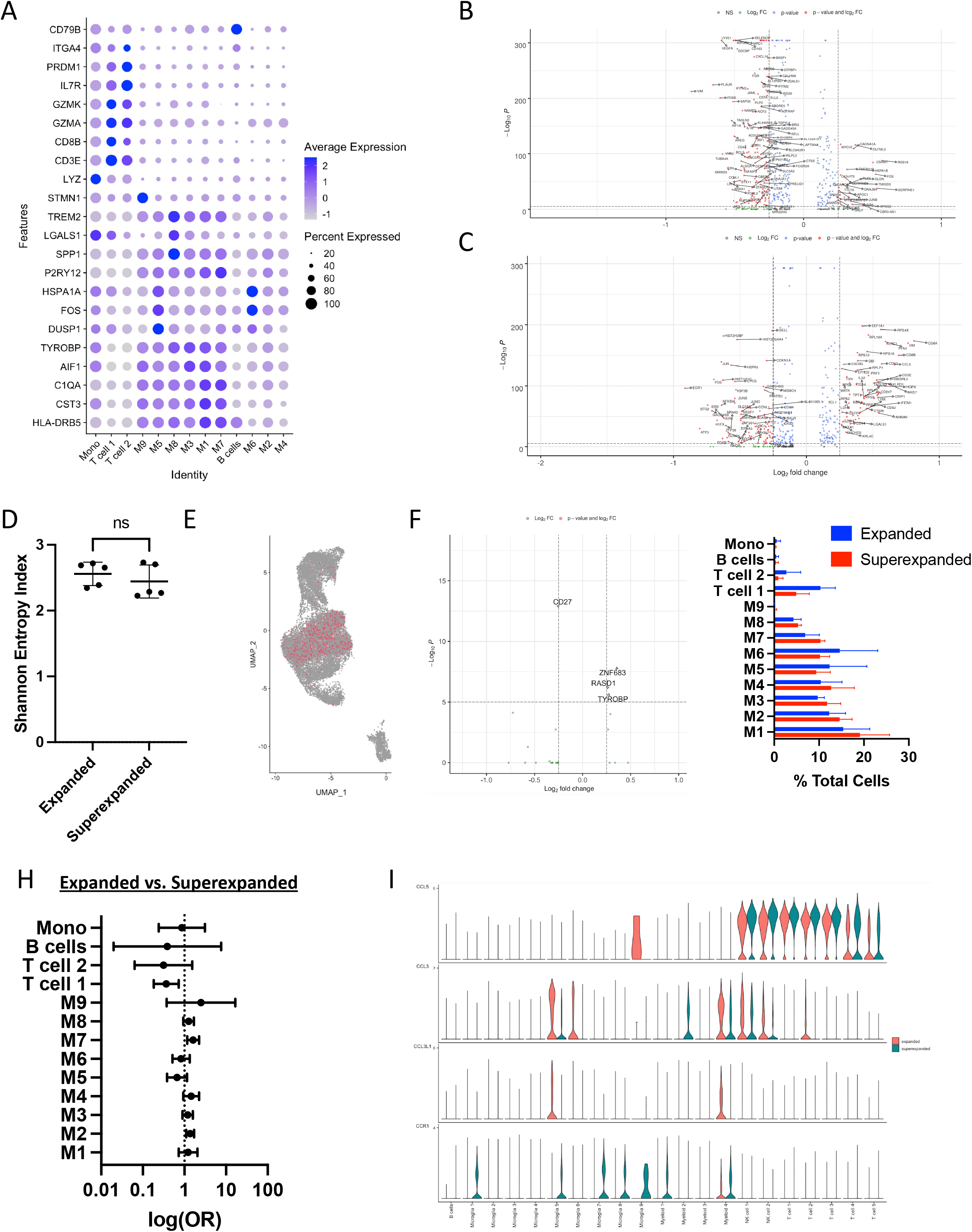
Meningeal T cells may regulate microglia immunity remotely via secreted factors. (A) DotPlot of brain immune cluster defining genes. Rows depict key marker genes, and columns represent distinct clusters. The blue color intensity represents the average expression of the corresponding marker in a cell cluster, while the size of the dot corresponds to the percentage of cells in that cluster that express the gene. (B) Volcano plot of genes differentially expressed between meningeal and brain myeloid cells. Positive Log2 fold change corresponds to genes upregulated in brain myeloid cells, and negative Log2 fold change corresponds to genes upregulate in meninges myeloid cells. (C) Volcano plot of genes differentially expressed between meningeal and brain T cells. Positive Log2 fold change corresponds to genes upregulated in brain T cells, and negative Log2 fold change corresponds to genes upregulated in meningeal T cells. (D) Shannon entropy index of brain TCR repertoires from patients with expanded and superexpanded meningeal TCR repertoires. t-test. (E) UMAP of meningeal lymphoid clusters with alpha/beta TCR+ cells with TCRs also found in the brain highlighted in red. (F) Volcano plot of genes differentially expressed between meningeal T cells with TCRs also found in the brain and all other CD8 T cells with a full alpha/beta TCR sequence. Positive Log2 fold change corresponds to genes upregulated in T cells with TCRs also found in the brain, and negative Log2 fold change corresponds to genes downregulated in T cells with TCRs also found in the brain. (G) Percentage composition of each brain immune cluster in samples with expanded and superexpanded meningeal TCR repertoires. (H) MASC analysis of brain cell cluster proportional changes between patients with expanded and superexpanded meninges. Odds ratio greater than 1 corresponds to an increased abundance of a particular cell cluster in the superexpanded group. Error bars correspond to 95% CI. (I) Violin plots of component ligands and receptors contributing to CellChat CCL signaling interaction compared between expanded and superexpanded meninges conditions.

